## Supplementary information for "The timing of differentiation and potency of CD8 effector function is set by RNA binding proteins"

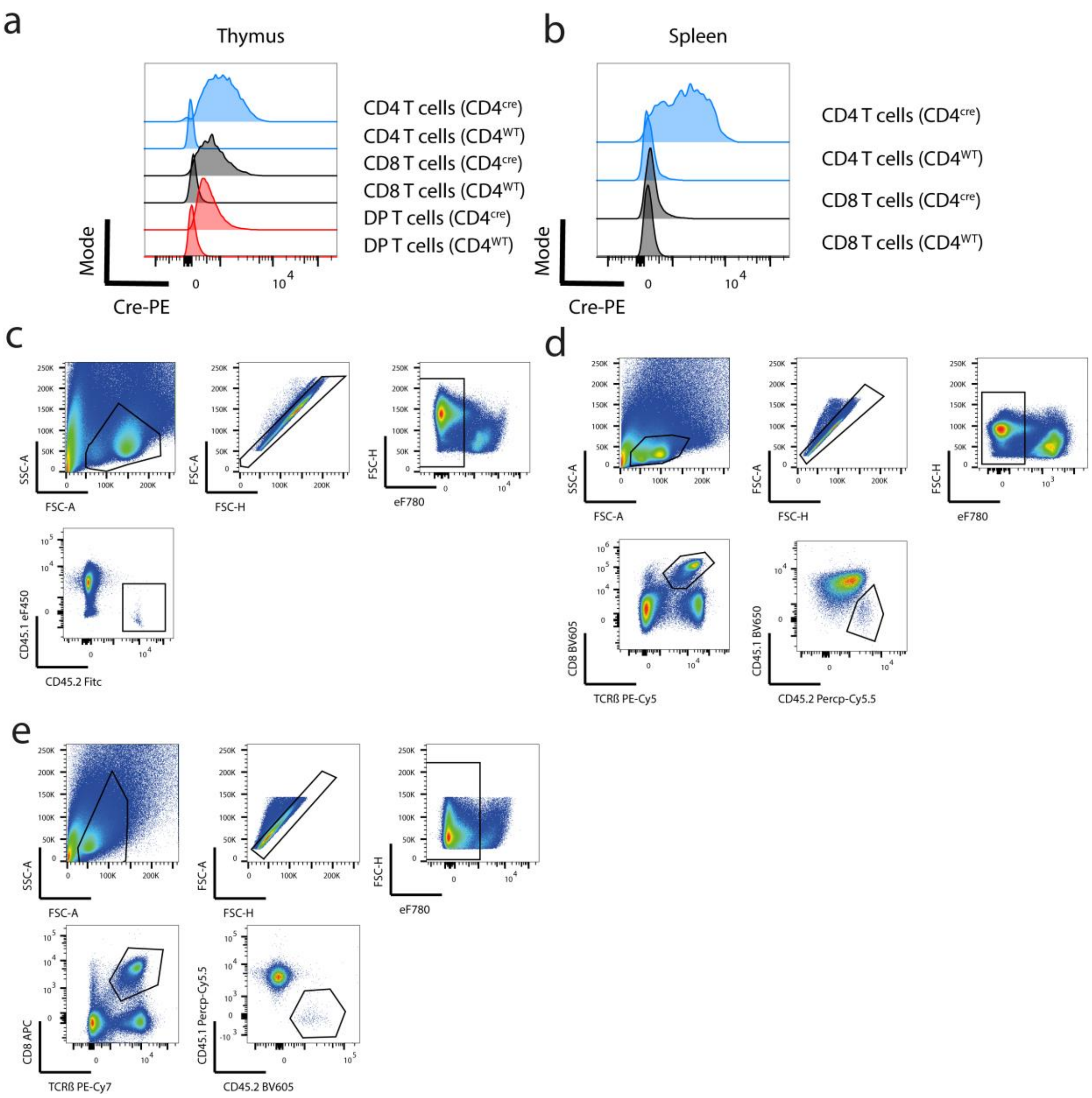


**Supplementary Figure 1. Cre expression in Thymocytes and Splenocytes and gating strategies.** Representative histograms showing expression of cre in single and douple positive CD4 and CD8 thymocytes (**a**) and mature CD4 and CD8 T cells (**b**) from OT-I transgenic CD4$^{cre}$ and CD4$^{WT}$ mice. **c-e)** gating strategies for analysis of transferred CD45.2 OT-I cells 10 days after influenza infections in lungs and spleens of CD45/1 recipient hosts.

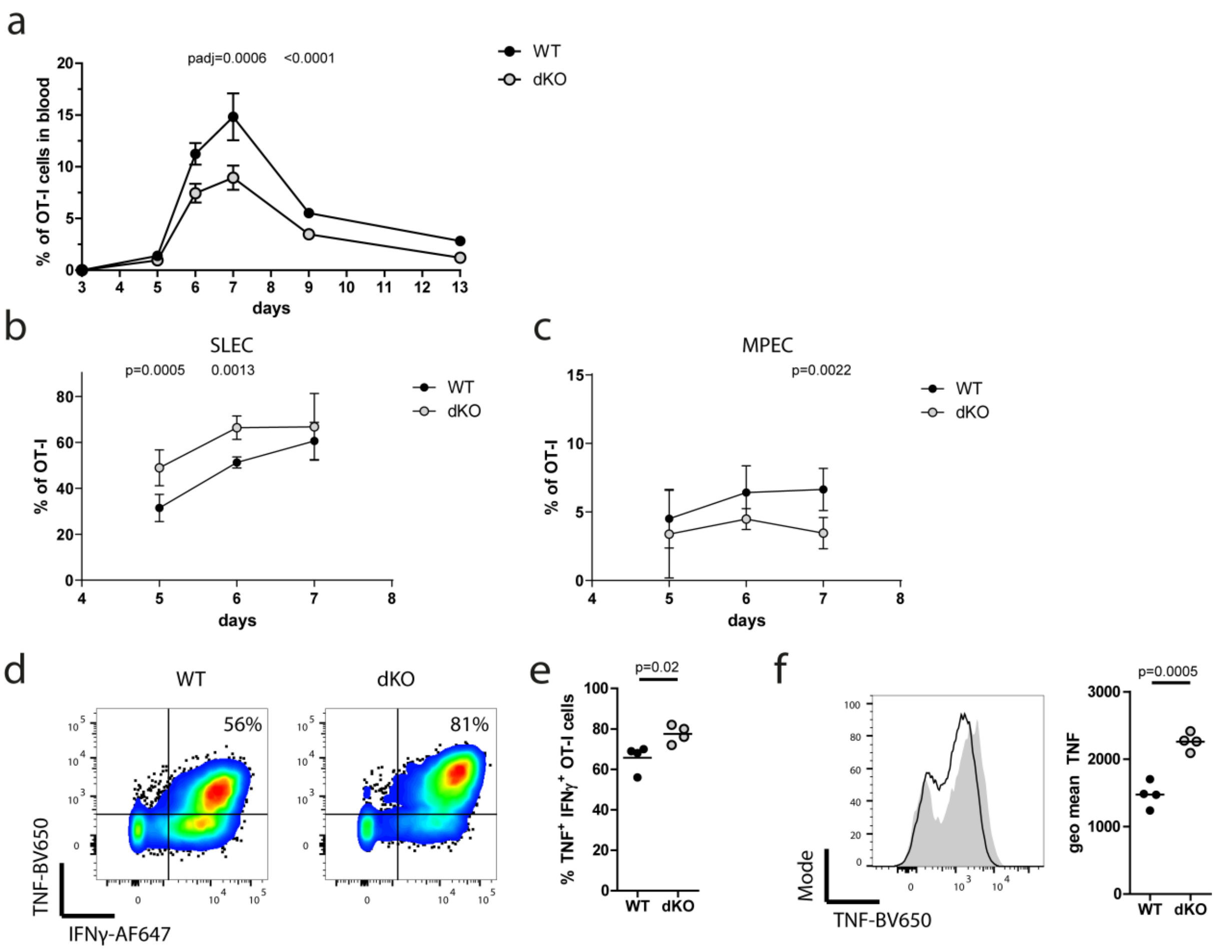


**Supplementary Figure 2. ZFP36 and ZFP36L1 limit rapid short lived effector CD8 T cell differentiation in response to *Listeria monocytogenes* infection. a)** Frequency of OT-I cells in the blood after infection with Lm-Ova. Data is compiled from two independent experiments. Error bars show the SEM. Statistical significance determined by two-way ANOVA followed by Bonferroni correction for multiple comparisons. Proportion of SLEC (**b**) and MPEC (**c**) over time after Lm-Ova infection. Data is pooled from two independent experiments and is presented as mean +/- SD. **d)** Representative plots of TNF and IFNγ staining in WT and dKO cells from blood on day 6 post infection following peptide restimulation (left panel). **e**) Frequency of TNF and IFNγ double positive effector cells. **f)** Representative overlay histogram of TNF expression in WT and dKO cells, the right panel shows the geometric mean fluorescence intensity of the TNF positive population. In b)-f) statistical significance was tested using a two-tailed unpaired Student's t-test. In all panels filled circles represent the respective WT control and open shaded circles dKO.

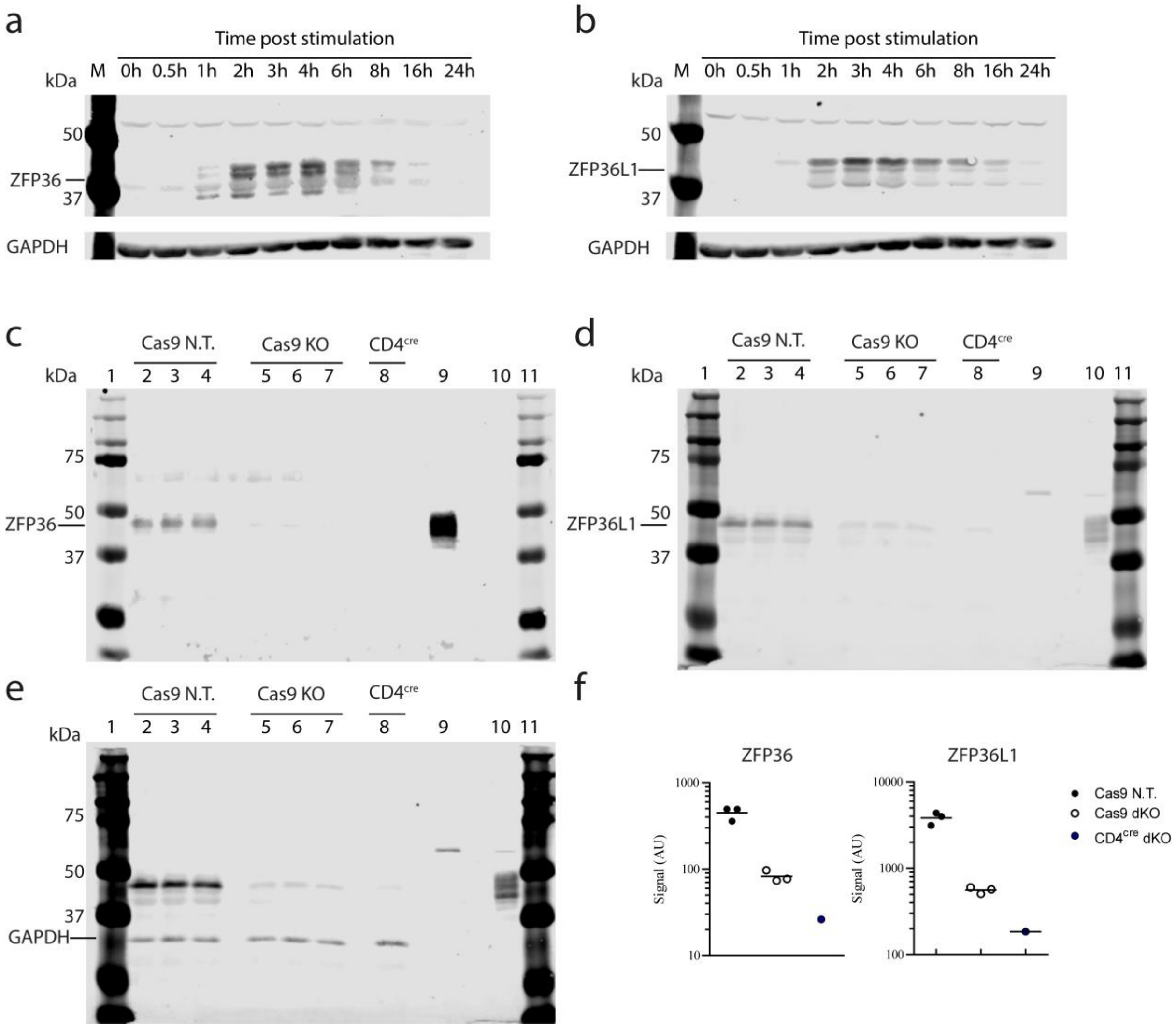


**Supplementary Figure 3. CrispR Cas9 mediated gene targeting of ZFP36 and ZFP36L1 leads to highly efficient ablation of both proteins**. (**a-b**) Western blot of ZFP36 and ZFP36L1 expression in CTLs stimulated with $10^{-7}$M N4 peptide for indicated times. Western blot analysis of **c)** ZFP36 **d)** ZFP36L1 and **e)** GAPDH in CTLs stimulated with $10^{-7}$M N4 peptide, where ZFP36 and ZFP36L1 were targeted for Cas9 mediated deletion. For all blots in c-e, the lanes were loaded as follows: 1: Protein marker; 2-4: lysates from three independently generated Cas9-GFP CTLs transduced with non-targeting guides; 5.-7: lysates from three independently generated Cas9-GFP CTLs transduced with guides targeting ZFP36 and ZFP36L1; 8: lysates from CTLs generated from CD4$^{cre}$ dKO mice. 9: lysate from HEK293 cells expressing ZFP36; 10: lysate from HEK293 cells expressing ZFP36L1; and 11: Protein marker. **f)** Quantification of ZFP36 and ZFP36L1 expression relative to GAPDH. Signal intensity is represented in arbitrary units (AU).

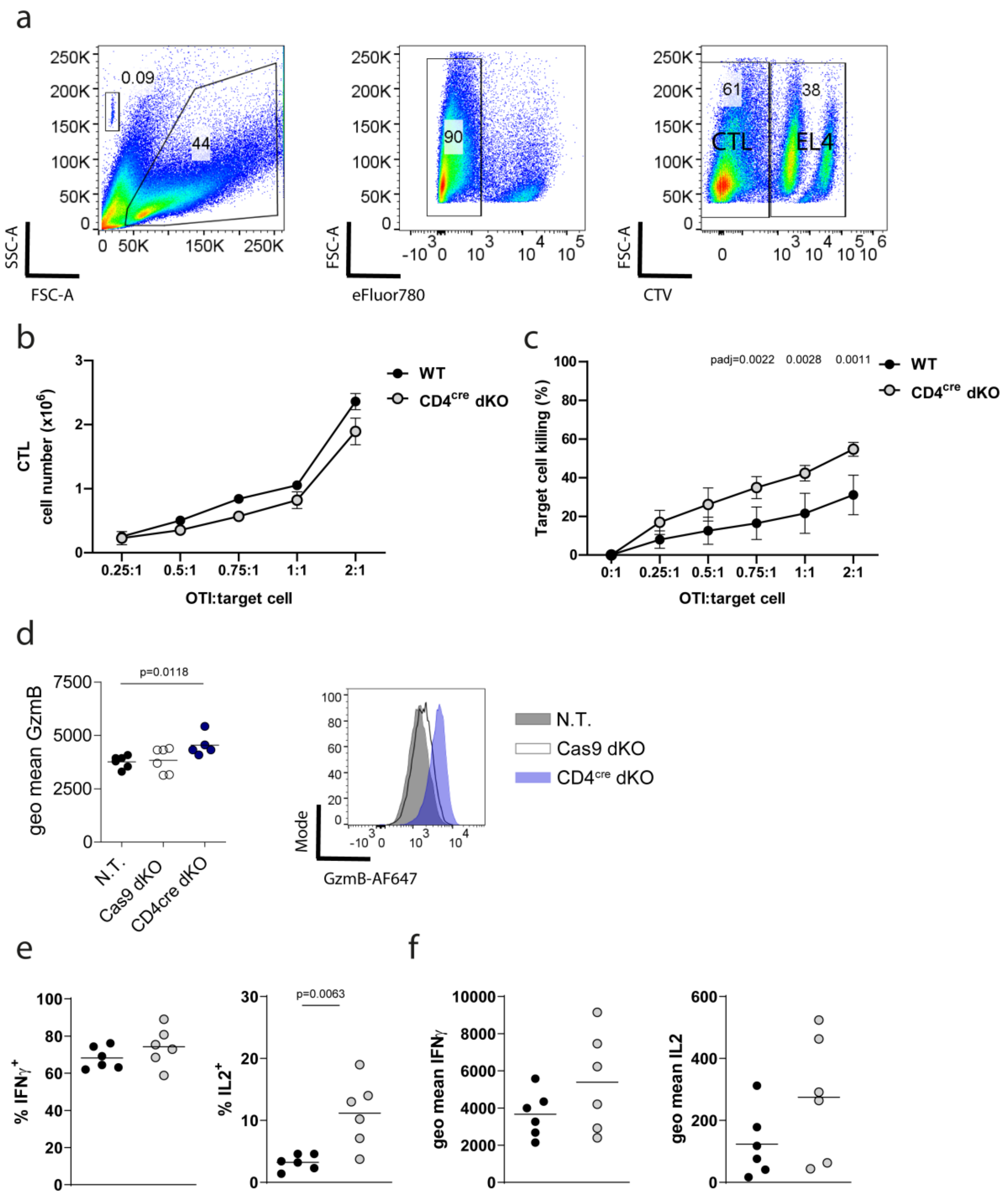


**Supplementary Figure 4. Zfp36 and Zfp36L1 deficient CTLs show enhanced killing potential in response to low affinity antigen.** (**a**) FACS plots represent CTLs, target and non-target EL4 cells seeded at a 1:1:1 ratio, following 3 hours of co-culture. Cells were pre-gated on FSC-A/SSC-A and live cells were determined as being eFlour780-. CTLs were defined as CTV-. Target EL4 and non-target cells were defined as CTV$^{hi}$ and CTV$^{lo}$, respectively. (**b**) CTL cell numbers recovered after 3h killing assay. Data is compiled from WT=2 and dKO=3 biological replicates. (**c**) In vitro killing of V4 peptide loaded EL4 cells after three hours at indicated CTL to EL4 ratios, using CTLs with CD4$^{cre}$ mediated deletion of

ZFP36 and ZFP36L1 and WT CTLs. Data is compiled from WT=8 and dKO=6 biological replicates. In b) and c) data is presented as mean values +/- SEM. Statistical significance was tested by two-way ANOVA analysis followed by Sidak's correction for multiple testing. (**d**) Geometric mean fluorescence intensity and representative FACS plots showing Granzyme-B expression in CTLs prior to antigenic stimulation (each point represents a biological replicate). Filled cricles and histograms show cells treated with non targeting guides and open circles and histograms show cells treated with RBP targeting guides. Blue open circle and histograms show cells derived from RBP dKO mice. **(e)** Frequencies and **(f)** geometric mean fluorescence of IFNγ and IL2 expression in WT and dKO CTLs stimulated with N4 peptide in the presence of Brefeldin A. In all panels if not otherwise stated filled circles represent the respective WT control and open shaded circles dKO; each dot represents a biological replicate. Statistical significance in panels d-f was tested using a two-tailed unpaired Student's t-test.

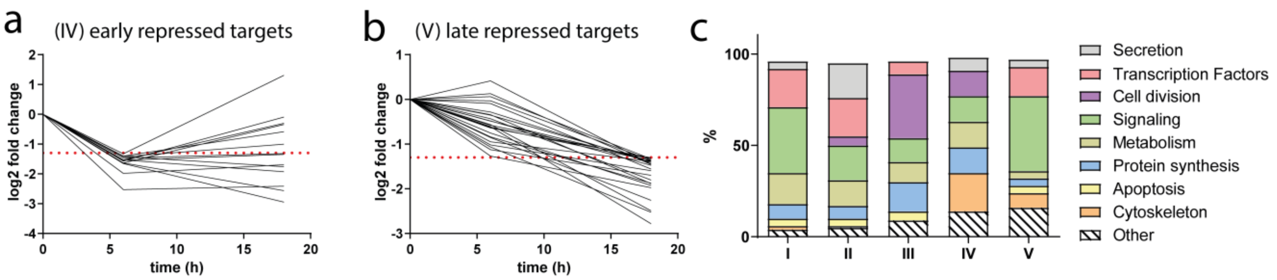


**Supplementary Figure 5. ZFP36 and L1 target transcripts are functionally distinct according to their temporal regulation**. **a, b)** Clusters IV and V showing ZFP36 family mRNA target dynamics during T-cell activation. **c)** Custom grouping of ZFP36 family targets according to function in cellular processes during T-cell activation. Dotted line represents the -1.3 log2 FC. Cluster I contains target genes which are induced at 6 hours but not at 18h. Cluster II contains target genes which are induced at both timepoints. Cluster III contains target genes which are induced only at 18h. Cluster IV and V contain target genes which are down regulated early and late respectively.

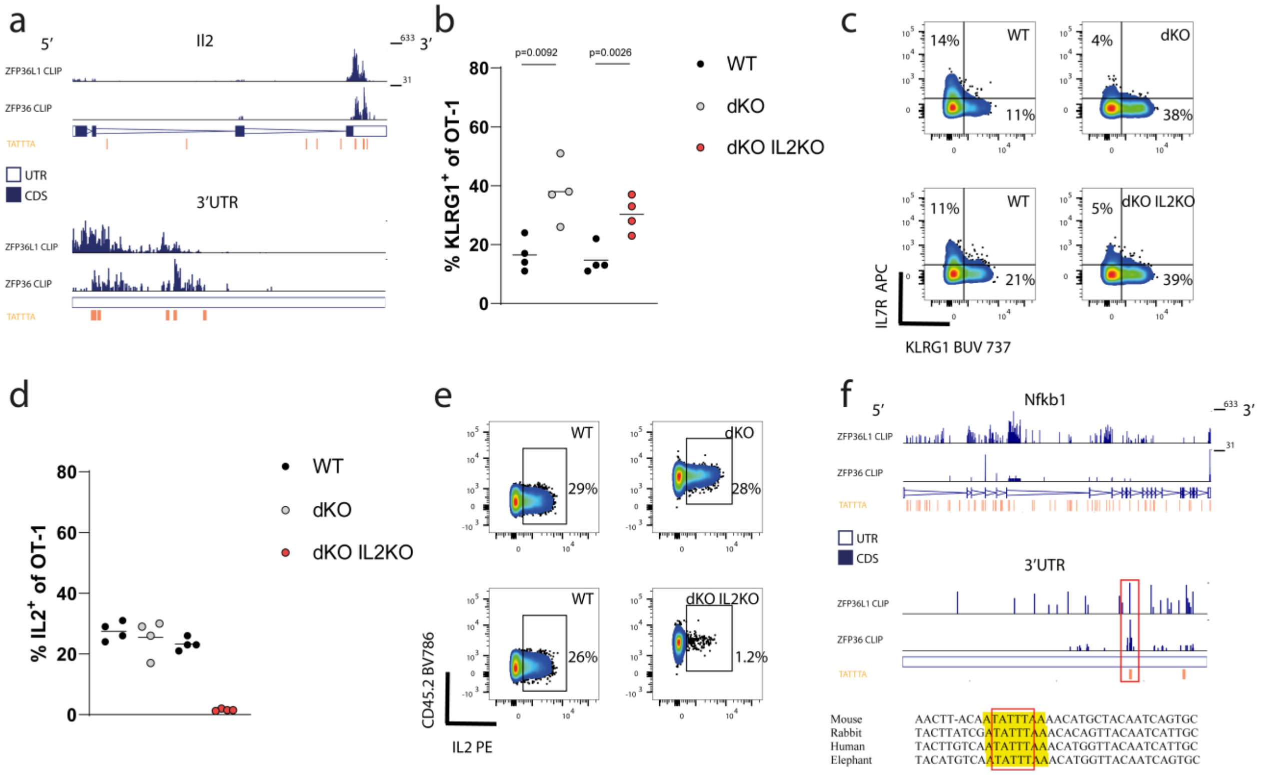


**Supplementary Figure 6. ZFP36 and ZFP36L1 directly target *Il2* and *Nfkb1*. a)** CLIP sequencing reads across the *Il2* transcript. Top set of two lanes depicts the distribution of reads over the whole gene. The bottom set of two lanes depicts the distribution of reads over the 3' UTR. In each set top the lane shows the reads for ZFP36L1 iCLIP data and the bottom lane shows the reads for ZFP36 HITS-CLIP data. **b)** Analysis of blood of mice transferred with 500 WT naive OT-I CD8 T cells (filled circle) which were cotransferred into CD45.1 recipient mice together with 500 OT-I dKO cells, which have been nucleoporated with Cas9 protein and control guides (open grey circle) or guides targeting IL2 (open red circle). Mice were analysed 5 days after infection with attLm-OVA, **c)** Representative FACS panels show the expression of IL7R and KLRG1 in cotrasferred OT-I cells (gated on CD8 and CD45.1 and CD45.2 respectively). **d)** Splenocytes of mice which received cotransferred OT-I cells were stimulated with N4 peptide on day7 post infection in the presence of BrefeldinA. Frequency of IL2 producing OT-I cells is shown. **e)** Representative flow cytometry plots show the expression of IL2 after N4 peptide restimulation. Statistical significance was determined using a two-tailed paired t-test. **f)** CLIP sequencing reads across the *Nfkb1* transcript (top set of the two lanes) and 3`UTR (bottom set of the two lanes). In each set the top lane shows ZFP36L1 iCLIP data and the middle lane shows the ZFP36 HITS-CLIP data. The bottom panel shows conservation among vertebrates of the ZFP36/ZFP36L1 binding site determined using the MULTIZ alignment in UCSC genome browser for *Nfkb1***.** Binding as identified by CLIP is highlighted in yellow. Selected species are ordered from top to bottom according the evolutionary proximity of the clades. Red bracket highlights the ARE. Source Data are provided as Source Data file.

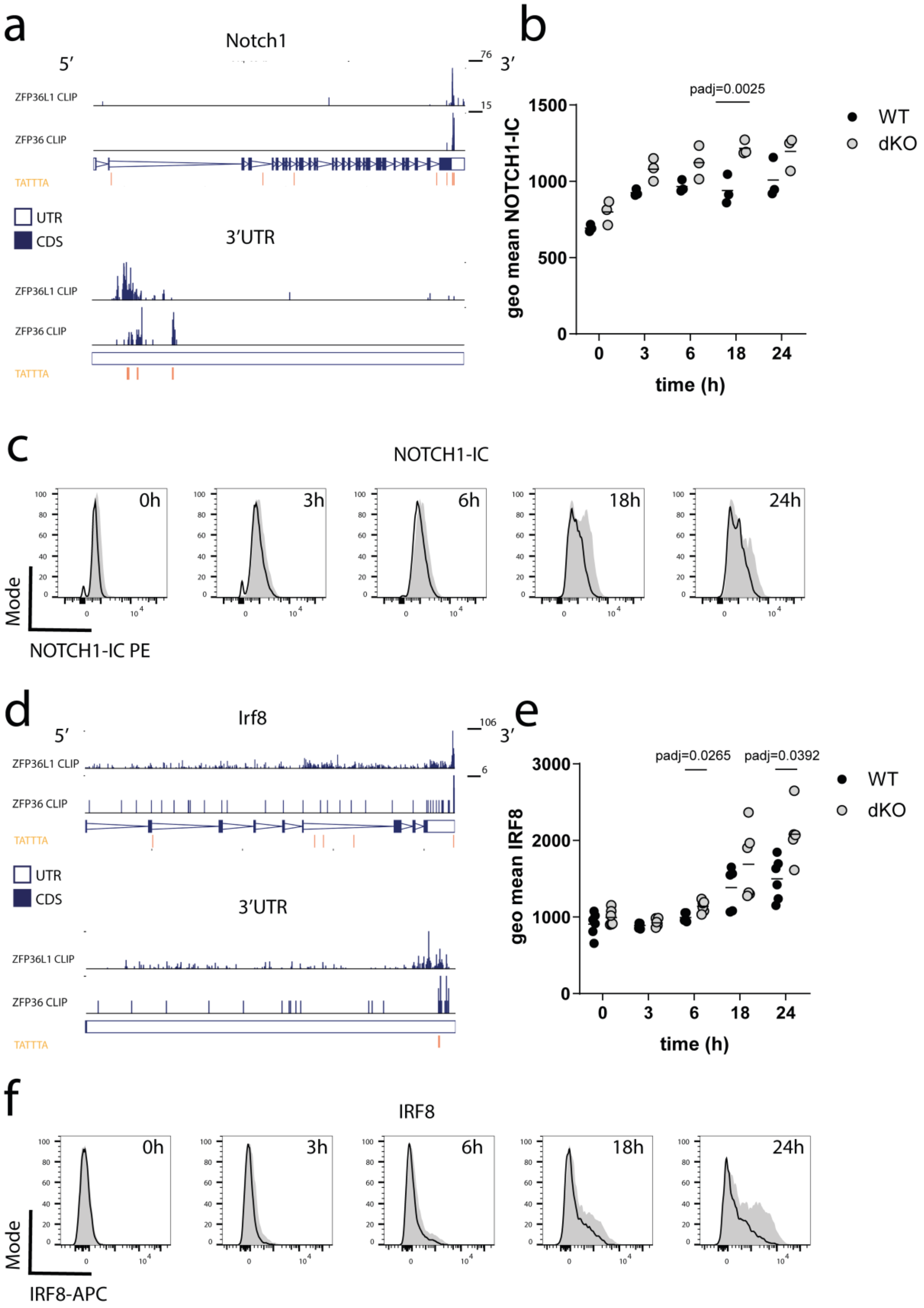


**Supplementary Figure 7: ZFP36 and ZFP36L1 directly target *Notch1* and *IRF8* to repress a transcription factor network driving effector differentiation. a)** CLIP data showing sequencing reads across the *Notch1* transcript (top set of the two lanes) with an expanded view of its 3'UTR (bottom set of the two lanes). In each set the top lane shows ZFP36L1 iCLIP data from OT-I CD8 CTLs stimulated for 3h with N4 peptide and the bottom lane shows ZFP36 CLIP data from in vitro activated naive CD4 T cells. The position of TATTTA motifs is shown. **b)** Summary of geometric mean fluorescence and **c)** representative flow cytometry for NOTCH1-IC. Open histograms represent WT and filled histograms show dKO cells. Statistical significance was tested by two-way ANOVA analysis followed by Sidak's test for multiple comparisons. Data is representative of three independent experiments. **d)** CLIP data showing sequencing reads across the *Irf8* transcript with an expanded view of its 3'UTR as described for *Notch1* in (a). **e)** Geometric mean of fluorescence of IRF8 expression by naive CD8 T cells following activation with plate bound anti-CD3 antibody. Statistical significance was determined using a mixed-effects model followed by Sidak's test for multiple comparisons. Data is compiled from 2 independent experiments. **f)** Representative flow cytometry histograms for IRF8. Open histograms represent WT and filled histograms show dKO cells. In all panels filled circles represent the respective WT control and open shaded circles dKO.

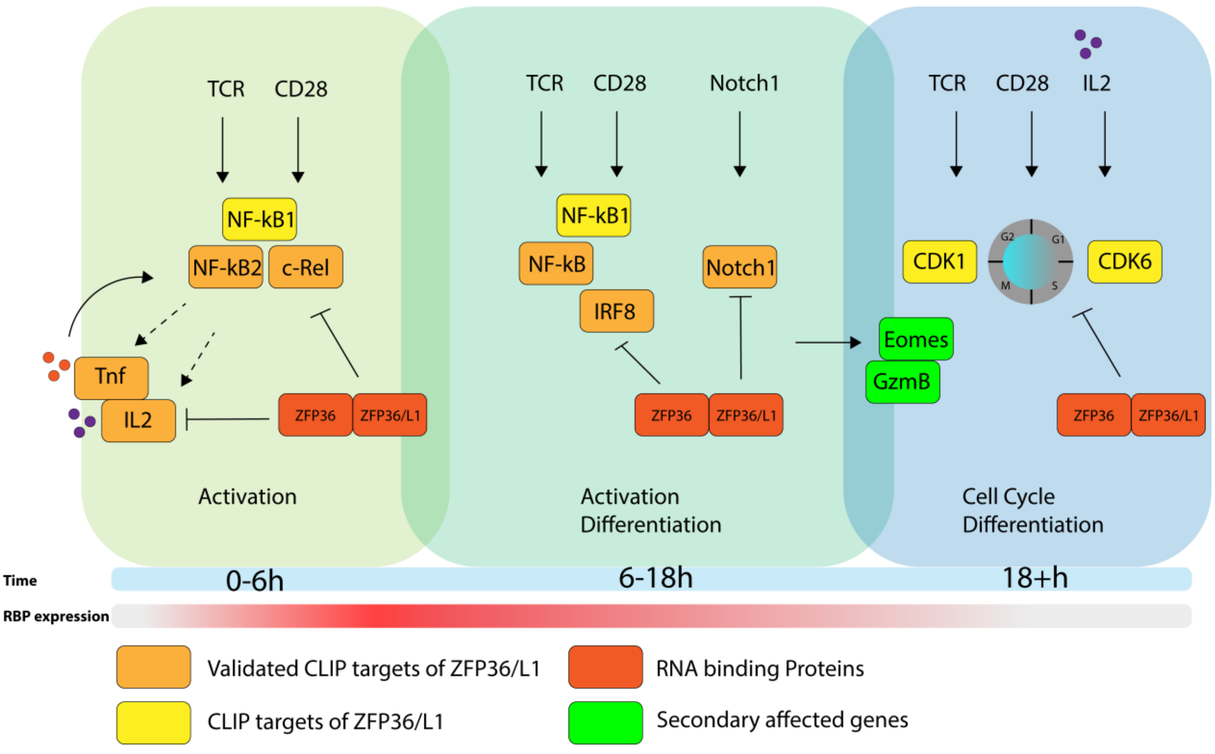


**Supplementary Figure 8. Model of ZFP36 and ZFP36L action in early T cell activation.** Schematic model of ZFP36 and ZFP6L1 mechanism of action during T cell activation.
